# Enhanced encephalitic tropism of bovine H5N1 compared to the Vietnam H5N1 isolate in mice

**DOI:** 10.1101/2024.11.19.624162

**Authors:** Kerry Goldin, Sarah van Tol, Randall C. Johnson, Reshma Koolaparambil Mukesh, Shane Gallogly, Jonathan E. Schulz, Jessica Prado-Smith, Greg Saturday, Kwe Claude Yinda, Vincent J. Munster, Emmie de Wit, Neeltje van Doremalen

## Abstract

In recent years, the landscape of highly pathogenic avian influenza (HPAI) virus infections has shifted, as evidenced by an increase in infections among mammals. This includes the recent circulation of H5N1 in dairy cattle herds in the USA and a rise in associated human cases. In this study, we investigated differences in tissue tropism of two HPAI H5N1 strains, the isolate A/Vietnam/1203/2004 (VN1203) isolated from a fatal human case in 2004 and the bovine isolate A/Bovine/Ohio/B24osu-342/2024 (Bov342) isolated in 2024, in C57BL/6J mice. Infection with either HPAI H5N1 isolate was uniformly lethal in mice. However, tissue tropism differed significantly: while VN1203 replication was largely restricted to the respiratory tract, Bov342 successfully replicated in the respiratory tract as well as various regions of the brain. Bov342-challenged animals exhibited clinical signs consistent with central nervous system (CNS) infection, and infectious virus was detected in brain tissue. Correspondingly, cytokine profiles in the brain differed significantly between the isolates. Notably, in addition to abundant evidence of CNS infection in Bov342-challenged mice via immunohistochemistry, sporadic intranuclear and intracytoplasmic immunoreactivity was observed in other tissues in the head, including the choroid plexus, retina, and inner ear. This study demonstrates that while both HPAI H5N1 isolates are uniformly lethal in C57BL/6J mice upon aerosol exposure, significant differences exist in tissue tropism, with Bov342 resulting in respiratory disease as well as increased neurotropism and inflammation in the brain and nasal turbinates compared to VN1203, which predominantly induces respiratory disease.

**Significance statement:** The rise in HPAI H5N1 infections among mammals, including humans in the USA, highlights an emerging One Health concern. Understanding the phenotypic changes of HPAI H5N1 and associated increase in infection of mammalian hosts is critical. In this study, we investigated the tissue tropism in mice of a bovine HPAI H5N1 strain isolated in 2024 and compared it to a strain isolated from a human patient in 2004. Our findings reveal that the bovine isolate exhibits enhanced neurotropism, unlike the respiratory-restricted replication observed with the HPAI H5N1 isolate from 2004. This difference in tissue tropism, accompanied by distinct cytokine responses in the brain, underscores the potential for altered disease outcomes in other mammalian hosts.

## Introduction

Recent years have seen an increase in spillover events where HPAI H5N1 viruses of clade 2.3.4.4b have crossed into a wide variety of species, including sea lions, mink, foxes, skunks, raccoons, and bobcats (1–3), which raises substantial concerns regarding the pandemic and panzootic potential of clade 2.3.4.4b HPAI H5N1 viruses. The increasing incidence of HPAI H5N1 virus in mammals underscores the urgent need for continued surveillance and research to better understand and mitigate the risk of a pandemic.

The first human case of HPAI H5N1 influenza virus infection was detected in 1997 in Hong Kong (4). Subsequently, HPAI H5N1 viruses rapidly diversified through reassortment with avian influenza viruses, leading to the emergence of a new genotype in clade 1: Z. Genotype Z spread throughout poultry in Southern China and was the only genotype detected by 2004 (5). Infection of poultry with genotype Z was also reported in China, Japan, South Korea, Thailand, Vietnam, Indonesia, Cambodia, and Laos (6). Human H5N1 infections were laboratory-confirmed in Vietnam, Cambodia, Indonesia, and Thailand, including fatal cases in Vietnam and Thailand (7, 8).

HPAI H5N1 (clade 2.3.4.4) was first detected in the United States and Canada in 2014 (6). A subclade, 2.3.4.4b, has displayed explosive expansion in wild birds (9) and was associated with fatal infections in mammals (1, 2, 10). Following reports of an abrupt drop in milk production and reduced feed intake by dairy cows in Texas and Kansas, milk and tissue samples from cattle tested positive for HPAI H5N1 by the Iowa State University Veterinary Diagnostic Laboratory (11). As of October 22, 2024, H5N1 has been reported in 492 herds across 15 states (12). Additionally, 25 human cases have been reported following exposure to dairy cows (13). A recent serologic survey across dairy workers demonstrated 7% (8 out of 115 persons) had evidence of a recent infection with H5N1 (14). One additional hospitalized case has not been associated with exposure to dairy cows or poultry, even though phylogenetic analysis shows clustering with bovine H5N1 isolates (15). All but one H5N1 human cases in the USA have been mild, resulting in conjunctivitis or respiratory symptoms (13).

In this study, we investigate whether the extensive evolution of HPAI H5N1 from genotype Z to the recent bovine HPAI H5N1 isolate results in differences in tissue tropism. We exposed C57BL/6J mice to either bovine H5N1 (A/Bovine/Ohio/B24osu-342/2024, Bov342) or HPAI H5N1 isolate obtained from the upper respiratory tract of a human case in Vietnam in 2004 (A/Vietnam/1203/2004 (VN1203), genotype Z) (16). Exposure to virus was done via aerosols, to mimic a more natural route of infection. We chose VN1203 as a comparison, since it has been extensively studied by other groups in animal models (17–20). Upon exposure, mice in both groups displayed reduced activity and rapidly lost weight.

Interestingly, whereas animals exposed to Bov342 showed neurological signs of disease, these signs were absent in animals exposed to VN1203. Accordingly, virus titers were high in brain tissue of Bov342- exposed animals compared to VN1203-exposed animals. Additionally, at endpoint cytokine levels were upregulated in brain tissue of Bov342-exposed animals, but not VN1203-exposed animals. Cytokine levels in lung tissues differed significantly between the isolates. Finally, immunohistochemistry showed markedly increased immunoreactivity in the brain, nasal turbinates and, to a lesser degree, the lungs in animals exposed to Bov342 compared to those exposed to VN1203. Thus, even though H5N1 exposure via aerosols is uniformly lethal for both VN1203 and Bov342 in C57BL/6J mice, tissue tropism differs, with Bov342 displaying a preference for the CNS and respiratory tract, whereas replication of VN1203 in the CNS is limited.

## Results

### C57BL/6J mice display dose-dependent signs of disease after aerosol challenge with HPAI H5N1

Mice were challenged with HPAI H5N1 A/Bovine/Ohio/B24osu-342/2024 (Bov342) or A/Vietnam/1203/2004 (VN1203) via aerosols, resulting in an inhaled dose of approximately 10^2^, 10^3^, or 10^4^ TCID_50_ (**Figure 1a**). Activity levels of six animals per group were monitored continuously and totaled per six hours. Compared to uninfected individuals, animals exposed to 10^4^ TCID_50_ Bov342 showed reduced activity starting at 48 hours post exposure. Consistent reduced activity was delayed by a further 18 hours in the 10^3^ dose group (66 hours post exposure) and 84 hours in the 10^2^ dose group (132 hours post exposure) compared to the 10^4^ dose group. Reduced activity in the animals exposed to VN1203 compared to uninfected animals was observed at 54 hours and 102 hours post exposure in the 10^4^ and 10^3^ dose group respectively but was not consistently observed in the 10^2^ dose group (**Figure 1b**). All six groups displayed weight loss compared to uninfected controls, although this did not reach significance in the 10^2^ dose VN1203 group (**Figure 1c**). Endpoint criteria were reached in all six animals within the Bov342 groups and 10^4^ and 10^3^ dose VN1203 groups, but two out of six animals did not reach endpoint criteria in the 10^2^ dose VN1203 group. Survival was dose-dependent; animals that received a lower dose of virus reached endpoint criteria later than those that received the highest dose of virus (**Figure 1d**). One of the two surviving animals was seronegative for neuraminidase (A/bald eagle/Florida/W22-134- OP/2022) and HA (VN1203) at 28 days post exposure, and thus did not have a productive virus infection (**Supplementary Figure 1a**-b). Clinical signs in mice differed depending on what virus isolate the animals were inoculated with: animals inoculated with Bov342 displayed signs indicative of infection of the central nervous system (CNS), including tremors and ataxia, whereas animals inoculated with VN1203 displayed primarily respiratory signs, including a hunched posture and tachypnea to dyspnea.

**Figure 1.**
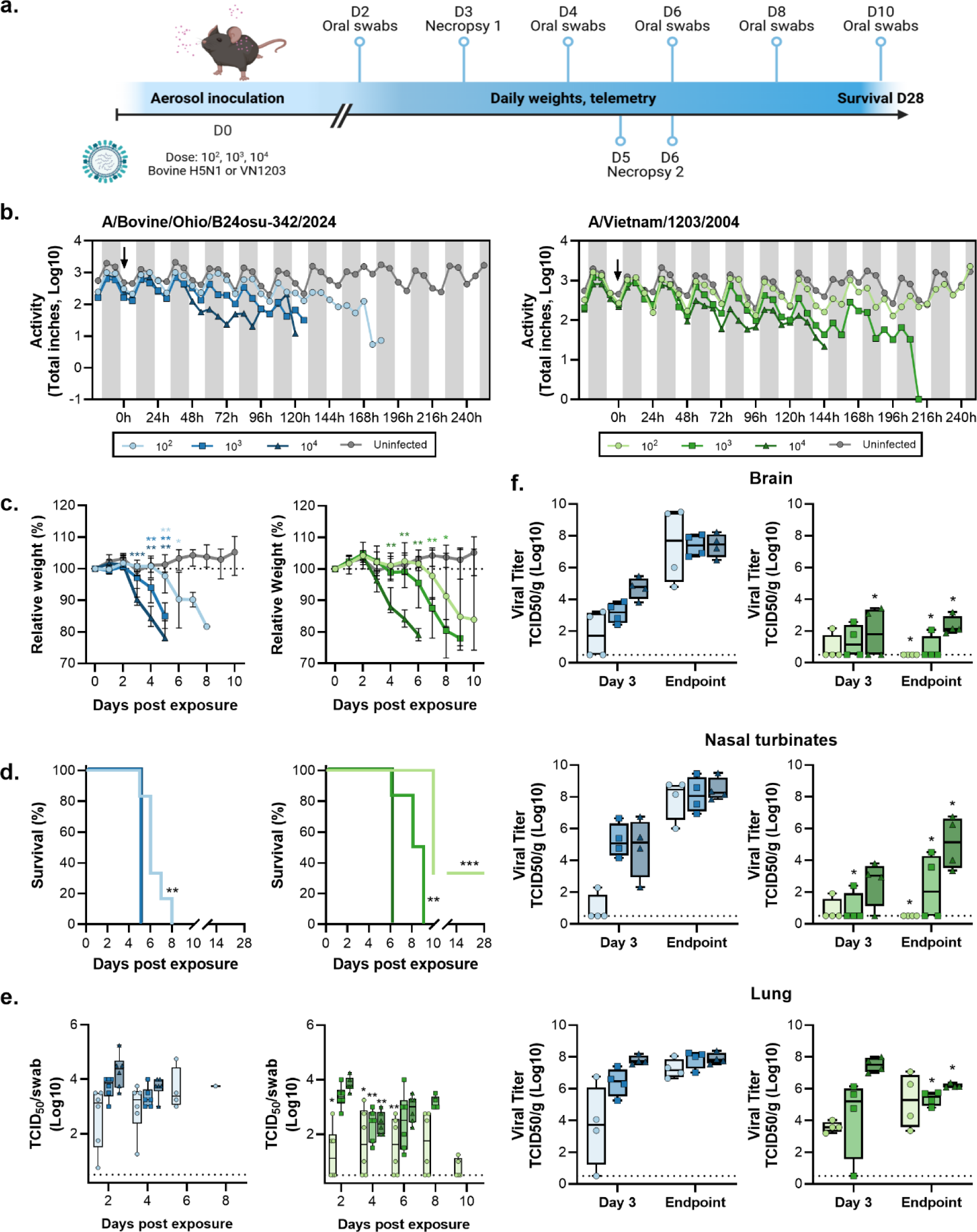
Aerosol inoculation of C57BL/6J mice with H5N1 results in dose-dependent mortality and virus replication. a. Experimental schedule. Mice (N=14) were inoculated with A/Bovine/Ohio/B24osu- 342/2024 or A/Vietnam/1203/2004 at 10^2^, 10^3^, or 10^4^ TCID_50_, via aerosols. Oral swabs were taken every two days (N=6), and necropsies were scheduled on D3 and D6 post exposure (N=4). One group per dose of virus was observed for survival up to D28 post exposure (N=6). Image created using Biorender. b. All animals in the survival groups received a transponder, and activity was monitored up to D10 post inoculation. Activity in inches traveled was totaled per 6 hours, shown is median (N=6). Arrow indicates time of virus exposure via aerosols. c. Relative weight compared to D0 post exposure. Shown is median with 95% confidence interval (CI), weights of all groups were compared to weights of control animals on same day post exposure. d. Survival post virus exposure (N=6). Survival comparison was done against high dose group per virus. e. Shedding profile of infectious virus in oral swabs. Individual values shown, bar graph shows minimum to maximum, with middle line displaying median (N=6). For each day, amount of shedding was compared between BoV342 and VN1203 animals that received the same dose of virus. f. Infectious virus titer in brain, nasal turbinate, and lung tissues at D3 and either D5 (BoV342) or D6 (VN1203) post exposure. Individual values shown, bar graph shows minimum to maximum, with middle line displaying median (N=4). For each day, amount of shedding was compared between BoV342 and VN1203 animals that received the same dose of virus. For statistical analysis in c, e, and f, first a one-way ANOVA (f), two-way ANOVA (c), or mixed-effect analysis (e) was performed. If significantly different, groups of interests were compared using a Mann-Whitney test. Survival was compared via Kaplan-Meier analysis. * = p-value <0.05; ** = p-value <0.01; *** = p-value <0.001.

Oropharyngeal swabs were taken at 2-, 4-, 6-, 8-, and 10-days post exposure. Infectious virus was detected in swabs obtained from all exposed animals, except for the two surviving animals in the 10^2^ dose VN1203 group. At 2-, 4-, and 6-days post exposure, virus shedding was higher in the 10^2^ dose Bov342 animals compared to the 10^2^ dose VN1203 animals. Significant differences in shedding between virus isolates for both the 10^3^ and 10^4^ dose groups were only observed on 4 days post exposure (**Figure 1e**). At 3 days post exposure, and when endpoint criteria were reached for at least one animal per virus (5 days post exposure for Bov342, and 6 days post exposure for VN1203), tissues were collected from 8 animals per group. Of four animals per group in the 10^4^ and 10^2^ dose groups only, relative lung weight was measured. Higher relative lung weight is indicative of fluid presence in the lung. At 3 days post exposure and at endpoint, a significant difference in relative lung weight was observed between the animals exposed to 10^2^ or 10^4^ TCID_50_ Bov342, whereas this was only observed at endpoint in animals inoculated with VN1203. No differences were observed between Bov342 and VN1203 when same dose inoculations were compared (**Supplementary Figure 1b**). Infectious virus titers were measured in brain, nasal turbinate, and lung tissues at 3 days post exposure and at endpoint. In line with the clinical signs that were observed, high viral titers were detected in brain tissue of animals that were inoculated with Bov342 (22 out of 24 animals tested positive). Viral titers were significantly lower in animals inoculated with 10^4^ dose VN1203 on day 3 and at endpoint, and for 10^2^ and 10^3^ dose VN1203 at endpoint (10 out of 24 animals tested positive). In contrast, infectious virus was detected in lung tissue of the majority of HPAI H5N1 exposed animals regardless of isolate (46 out of 48 animals). Significant differences in virus titer in lung tissue between virus isolates were noted in 10^3^ and 10^4^ dose groups at endpoint, but not at 3 days post inoculation. Infection of nasal turbinates mirrored that of brain tissue; 21 out of 24 tissues of animals inoculated with Bov342 were positive, whereas 11 out of 24 tissues of animals inoculated with VN1203 tested positive, and titers were higher in animals exposed to Bov342 (**Figure 1f**).

### C57BL/6J mice inoculated with HPAI H5N1 exhibited differential cytokine and chemokine responses corresponding with virus isolate, time of necropsy, and inoculum dose

We then investigated cytokine and chemokine expression in brain, nasal turbinate, and lung tissues obtained from the 10^2^ and 10^4^ dose groups. In the brain (**Figure 2a and Supplementary Figure 2a**), infection with Bov342 induced the expression of type-I (*Ifna4*, *Ifnb*), -II (*Ifng*), and -III (*Ifnl2*) interferons (IFN), interferon stimulated genes (ISGs) (*Isg15*, *Ifit1*, *Mx1*, *Oas1*), and pro-inflammatory cytokines (*Il1b*, *Il6*, and *Tnfa*) at endpoint. The magnitude of induction was increased at endpoint compared to day 3 for *Ifna4*, *Ifng*, *Il6*, *Tnfa*, and ISGs, and expression was dose-dependent for *Ifna4*, *Infg*, and *Tnfa*. With the exception of *Ifna4*, the evaluated genes were not induced in the brain of mice infected with VN1203. Further, at endpoint, expression of *Ifna4*, *Ifng*, *Il1b*, *Il6*, *Tnfa*, and ISGs were significantly higher in mice infected with Bov342 than VN1203.

**Figure 2.**
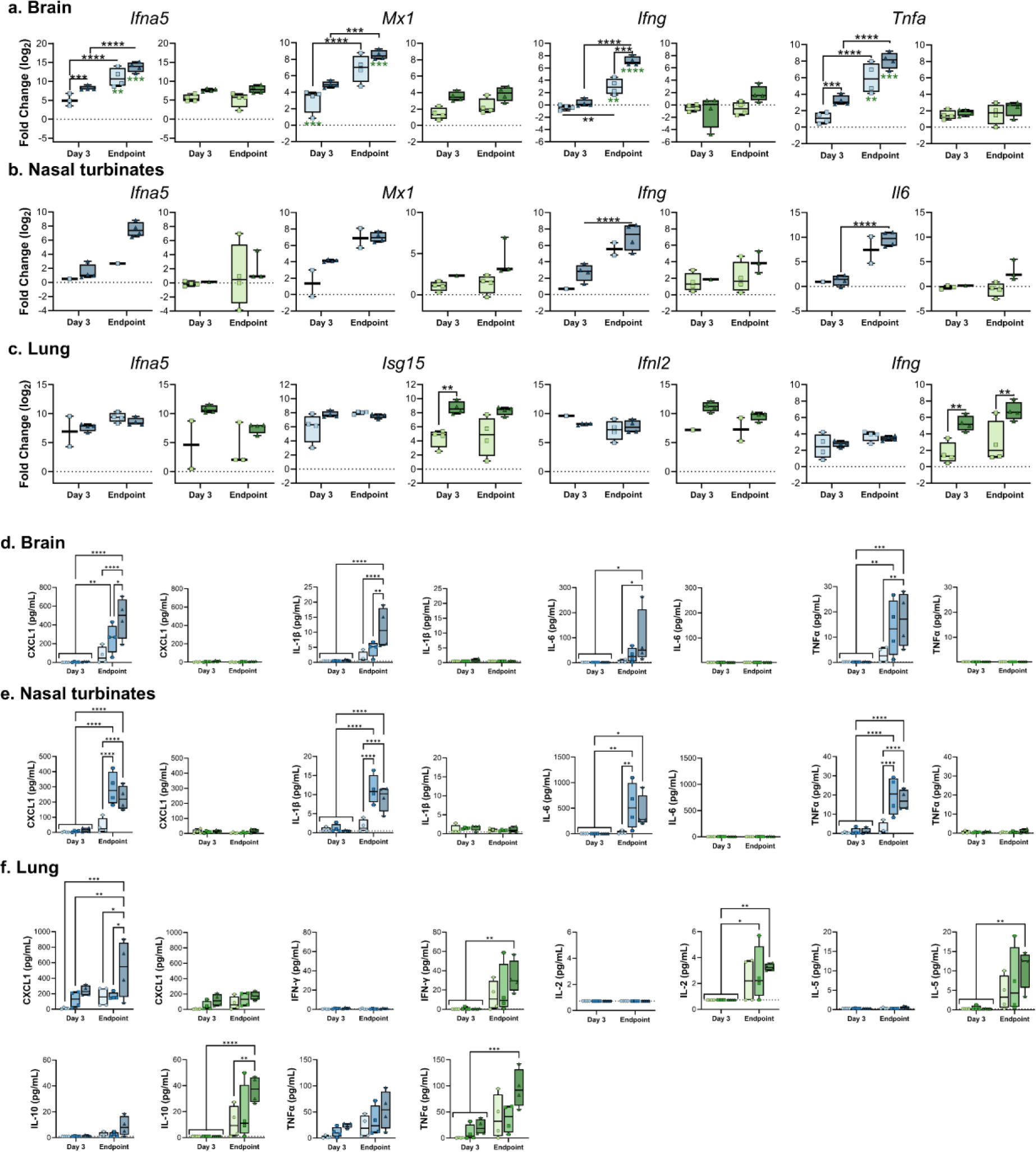
Aerosol inoculation of C57Bl/6J mice with HPAI H5N1 results in virus-, tissue- and dose- dependent induction of cytokines and antiviral effectors. Gene expression in brain (a), nasal turbinate (b), or lung (c) shown as log_2_ transformed fold change relative to healthy control mice. Cytokine and chemokine detection in brain (d), nasal turbinate (e), or lung (f) tissue. Individual values shown, bar graph shows minimum to maximum, with middle line displaying median (N=4). Gene expression differences between viruses and dose were compared at day 3 and endpoint using a two-way ANOVA with Tukey’s post-test, and differences across time points were assess with a two-way ANOVA with Sidak’s multiple comparisons test. A Bonferroni correction was applied, and only p-values greater than 0.0044 were considered significant. **= p-value <0.0044, ***= p-value <0.001, ****=p-value <0.0001.

The gene expression changes in nasal turbinates mirrored those observed in the brain. Infection with 10^4^ dose Bov342 induced type-I, -II, and -III IFN, ISGs, and pro-inflammatory cytokines at the endpoint, but infection with either 10^2^ or 10^4^ dose of VN1203 did not induce differential expression of any genes assessed (**Figure 2b and Supplementary figure 2b**). Due to insufficient nasal turbinate sample for one of the 10^2^ dose Bov342-exposed mice and variability within its endpoint group, we were unable to statistically determine dose-dependent differences.

Within the lungs of mice challenged with 10^4^ TCID_50_ VN1203, type-I, -II, and -III IFN, ISGs, and pro-inflammatory cytokines genes were induced at day 3 and stayed elevated or decreased in expression at endpoint (**Figure 2c and Supplementary Figure 2c**). Exposure to 10^2^ dose VN1203 did not induce expression of any genes uniformly at either timepoint, and induction of *Ifng, Isg15*, and *Tnfa* were significantly lower than mice challenged with 10^4^ TCID_50_ VN1203. Virus-dependent differences in gene expression were less apparent in the lung than in the other tissues. Although none of the differences were statistically significant, *Ifng* expression in mice infected with Bov342 tended to be lower than those with VN1203.

Protein levels of cytokines and chemokines were then assessed in brain, nasal turbinate, and lung tissue across all dose groups (**Figure 2 and Supplementary** Figure 2). In both brain and nasal turbinate tissues from animals exposed to Bov342, but not VN1203, we detected a dose-dependent expression of CXCL1, IL-1β, IL-6, and TNF-α at endpoint.

In lung tissue of Bov342-exposed animals, a dose-dependent elevation of CXCL1 and TNF-α was observed, particularly in tissues collected at endpoint. In contrast, elevated levels of IFN-γ, IL-2, IL-5, IL- 10, and TNF-α were detected in lung tissue obtained from animals exposed to VN1203 at endpoint.

We then performed Principal Component Analysis (PCA) of viral titer, protein abundance, and mRNA levels in the brain, lung, and nasal turbinates of mice, separated by day of harvest (**Figure 3**). Analysis at 3 days post inoculation resulted in two distinct clusters: the smaller cluster contained only lung samples, all but one of which were high dose groups. The remaining low-dose lung samples, all brain, and all nasal samples were contained in the larger cluster.

**Figure 3.**
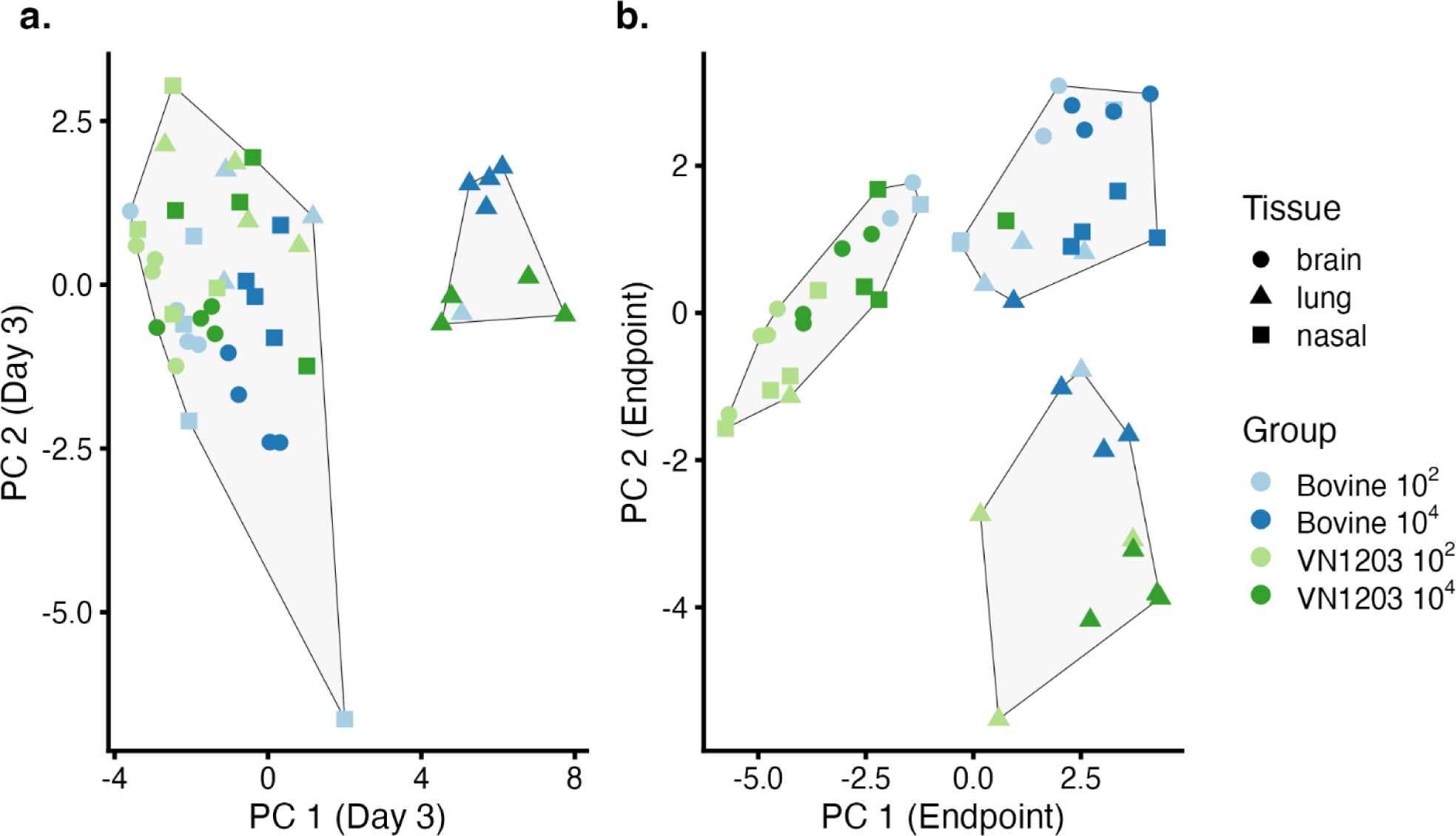
Principal Component Analysis of virus titer, protein abundances, and mRNA levels in mice grouped by tissue, virus, and dose. Principal component analyses are shown for mice at 3 days post inoculation (a) and at endpoint (b) with K-means clusters differentiated by light grey polygons. The optimal number of clusters (two for day 3 and three for endpoint) were identified by the majority vote of 26 indices for determining the number of clusters (1). 1. M. Charrad, N. Ghazzali, V. Boiteau, A. Niknafs, NbClust: An R Package for Determining the Relevant Number of Clusters in a Data Set. *Journal of Statistical Software* **61**, 1 - 36 (2014).

In the PCA of samples collected at endpoint, the optimal number of clusters was determined to be three, with clusters similar in size. One cluster was primarily composed of VN1203-exposed samples, while a second cluster mainly contained Bov342-exposed samples, both predominantly from brain and nasal turbinate samples. The third cluster contained only lung samples, where all but one of the VN1203- exposed mice clustered together with most of the high-dose Bov342-exposed mice. Most of the low-dose Bov342-exposed lung tissues clustered with the Bov342-exposed brain and nasal samples.

### Histopathologic lung lesions are similar in C57BL/6J mice exposed to both H5N1 isolates and influenza A NP immunoreactivity distribution is virus-dependent

Histologically, pathology was primarily observed in the lungs in both Bov342 and VN1203-exposed animals (**Figure 4a**). At 3 days post exposure, animals exposed to the 10^4^ dose of both isolates had developed pulmonary lesions, with lesions being more severe in the Bov342-exposed animals (**Figure 4a**). At endpoint, for the 10^4^ dose for both isolates, histologic lesions were similar to those observed at the 3 days post exposure timepoint **(Figure 4a**). Animals exposed to 10^2^ TCID_50_ of Bov342 did not have any pulmonary lesions at 3 days post exposure or endpoint (**Figure 4a**). VN1203-exposed animals in the 10^2^ dose group did not have pulmonary lesions at 3 days post exposure, however, at endpoint two of the four animals had developed moderate pulmonary lesions (**Figure 4a**). The most common finding in all groups was an acute, moderate to severe, multifocal, necrotizing bronchiolitis, characterized by erosion and ulceration of the bronchiolar epithelium and abundant degenerate cellular debris and fibrin within the bronchiolar lumen, occasionally forming occlusive plugs (**Supplementary Figure 3a**). Multifocal, moderate, bronchointerstitial pneumonia was also observed, characterized by lymphoplasmacytic infiltrates within the peribronchiolar space and expanding the surrounding alveolar septa (**Supplementary Figure 3a**). On immunohistochemistry for influenza A virus NP in the lungs, in both Bov342 and VN1203 animals in the 10^4^ dose groups, animals were given a moderate semiquantitative score at 3 days post exposure and a higher score at endpoint (**Figure 4b**). In the 10^2^ dose Bov342 animals, two of the four animals had mild to moderate immunoreactivity at 3 days post exposure, and at endpoint one of four animals had marked IHC positivity. In the 10^2^ dose VN1203 group, only one animal from both 3 days post exposure and endpoint displayed moderate IHC positivity (**Figure 4b**). Intranuclear and intracytoplasmic immunoreactivity in all groups was primarily within bronchiolar epithelium, and necrotic debris within airway lumen (**Figure 4c**). Immunoreactivity was observed in the nasal respiratory epithelium primarily in the 10^4^ dose Bov342 animals at both 3 days post exposure and at endpoint (**Figure 4d**). Immunoreactivity also extended into alveoli, with both type I and type II pneumocytes being infected (**Figure 4c**). Immunoreactivity in pneumocytes was typically not associated with observed pathology (**Figure 4c**).

**Figure 4.**
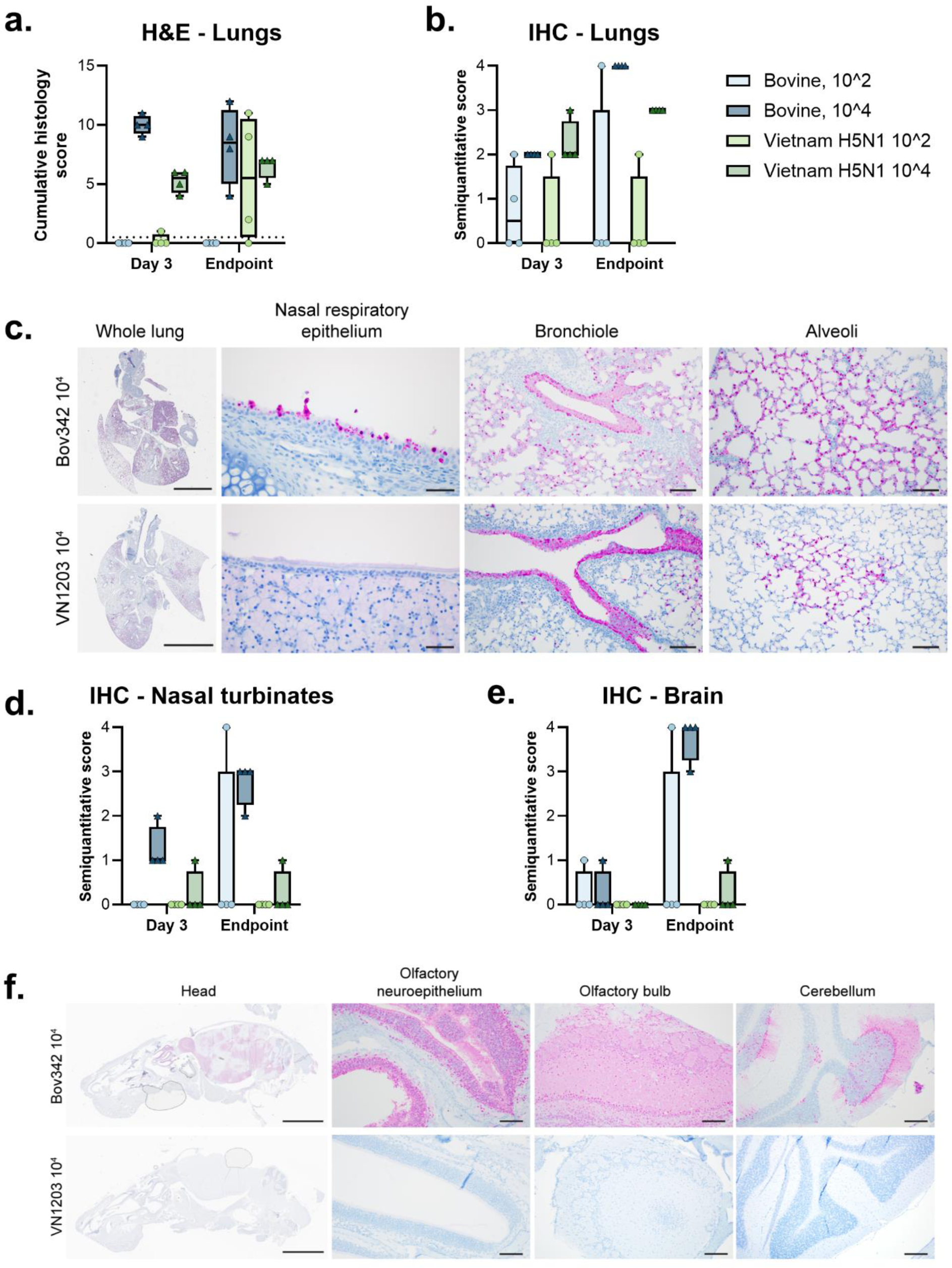
Aerosol inoculation of C57BL/6J mice with H5N1 results in similar histopathologic lesions in the lungs and virus-dependent IAV NP immunoreactivity in the olfactory epithelium and CNS. . Pulmonary lesions were observed in the Bov342 10^4^ group and both VN1203 doses at 3 days post exposure and at endpoint (a). Cumulative histology score calculated by adding semiquantitative values assigned in multiple categories (Table S2) (a). Immunoreactivity for influenza A virus NP (pink) was observed in the lungs in all animals in the 10^4^ dose groups for animals at 3 days post exposure and endpoint, however, only observed sporadically in the 10^2^ dose groups for both isolates at both timepoints (b). At endpoint, immunoreactivity in the Bov342 10^4^ group was observed within the nasal respiratory epithelium and lower airways, including bronchioles and type I and type II pneumocytes. Comparatively, VN1203-exposed animals had similar immunoreactivity in the bronchioles and alveoli and rare positivity in the nasal respiratory epithelium at endpoint (c). Influenza A virus NP positivity was observed in the nasal turbinates of the Bov342 10^4^ group at both timepoints, in one Bov342 10^2^ animal at endpoint, and in one animal in the VN1203 10^4^ group and 3 days post exposure and endpoint (d). At endpoint, immunoreactivity in the brain was observed abundantly in all Bov342 10^4^ animals and one animal in the Bov342 10^2^ group (e). Immunoreactivity in the Bov342 10^4^ animals at endpoint was widely distributed throughout the olfactory neuroepithelium and CNS, including the olfactory bulb and randomly scattered sites throughout the cerebrum, cerebellum, and brainstem (f). VN1203-exposed animals had no significant positivity in the olfactory tissue or CNS (f). For all graphs, individual values shown on boxplots. Whole lung: scale bar=4mm. Nasal respiratory epithelium: scale bar=50µm. Bronchiole: scale bar=100µm. Alveoli: scale bar=100µm Head: scale bar=5mm. Olfactory neuroepithelium: scale bar=100µm. Olfactory bulb: scale bar=200µm. Cerebellum: scale bar=200 µm.

Histologic lesions were not observed for animals exposed to either isolate in nasal turbinates or the brain with the exception of one animal in the 10^2^ dose Bov342 group at 3 days post exposure, where very mild acute lesions were observed in the olfactory neuroepithelium and olfactory bulb **(Supplementary Figure 3b**). The olfactory neuroepithelium in the 10^4^ dose Bov342 animals was moderately positive on IHC (**Figure 4d, 4f**). In the brain, immunoreactivity was marked in the 10^4^ dose Bov342 animals at endpoint, with all regions of the brain being affected despite the complete lack of pathologic lesions (**Figure 4e, 4f**). Immunoreactivity in the nasal turbinates and brain of the 10^2^ dose Bov342 group and all VN1203 groups was rare (**Figure 4d, 4e**). Interestingly, immunoreactivity was observed in other tissues in the head, which were occasionally captured in the sagittal sections. In multiple 10^4^ dose Bov342 animals, influenza A virus NP antigen was sporadically observed in the choroid plexus and ependymal cells lining the ventricle (**Supplementary Figure 4a**), cranial nerve nuclei along the ventral aspect of the calvarium (**Supplementary Figure 4b**), multiple cell layers of the retina (**Supplementary Figure 4c**), skeletal myocytes (**Supplementary Figure 4d**), mucosal epithelium overlying the hard palate (**Supplementary Figure 4e**), and within brown adipocytes (**Supplementary Figure 4f**). In a single 10^4^ dose Bov342 animal, immunoreactivity was observed in the dental pulp and overlying odontoblasts of a tooth root (**Supplementary Figure 4g**). In one 10^2^ dose VN1203 animal, epithelial cells lining the inner ear were immunoreactive (**Supplementary Figure 4h**). Importantly, the immunoreactivity in all these sites was both intranuclear and intracytoplasmic, indicating virus replication in these tissues.

## Discussion

In the current manuscript, we show that exposure to HPAI H5N1-containing aerosols results in productive virus replication in C57BL/6J mice, regardless of whether animals were exposed to HPAI H5N1 strain VN1203 or bovine isolate Bov342. We utilized three different exposure doses, ranging from approximately 10^2^ to 10^4^ per animal. Importantly, all animals in which evidence of H5N1 virus replication was detected reached endpoint criteria; only two animals in the low-dose VN1203 group did not reach endpoint criteria and did not seroconvert, suggesting they were likely not infected upon exposure. This is supported by the histologic findings, wherein only two of the four 10^2^ dose VN1203 animals developed significant pulmonary lesions. This study is not powered to define a 50% infectious dose (ID50), but based on these results, it is likely that the ID50 of VN1203 via aerosol exposure in C57BL/6J mice is around 10^2^ TCID_50_. Additionally, it appears that productive infection is 100% lethal in this model.

One out of two survivors did not show any evidence of seropositivity. This animal was co-housed with the second survivor (which was seropositive for NA, but not HA) and a third H5N1 VN1203-exposed mouse for the entirety of the experiment. Although this third animal reached endpoint criteria at 10 days post exposure and infectious virus was detected in three out of five oropharyngeal swabs, no evidence of H5N1 transmission to the seronegative mouse was detected. This observation aligns with a recent study, in which no transmission of a bovine isolate of H5N1 was observed between adult mice, whereas transmission from infected mother to pups was detected (18).

The most striking finding in this study was the differing tissue tropism between animals exposed to VN1203 and Bov342. Only Bov342-exposed animals exhibited productive virus replication in brain and nasal turbinate tissue, accompanied by signs of CNS infection and distinct cytokine and chemokine profiles. A recent study involving the inoculation of 6-week-old C57BL/6J mice with VN1203 and Bov342 via intranasal and oral routes also reported a CNS tropism for Bov342, but not for VN1203 (21). This contrasts with other studies where VN1203 was found to infect brain and nasal turbinate tissue (17–19). In two of these studies, BALB/cJ mice were inoculated via the intranasal route, whereas in our study C57BL/6J mice were exposed via aerosols, and therefore, the observed tropism may be mouse strain dependent or inoculation route dependent. It is well known that different inoculation routes will alter where the virus starts initial replication (22–24). Interestingly, in a SARS-CoV-2 mouse model in which the human receptor is overexpressed in all tissues, exposure via aerosols results in respiratory disease but no neurological invasion, like we observe with VN1203. In contrast, inoculation via the intranasal route results in both respiratory and neurological disease (25). Contrastingly, in an inoculation route comparison study in which BALB/c mice were inoculated with either HPAI H5N1, LPAI H7N9, or 2009 H1N1 via aerosol exposure or direct intranasal inoculation, no differences in lethality were observed based on inoculation route (26). Notably, bovine H5N1 was not neurotropic in BALB/c mice and no neurological signs of disease were described, in contrast to bovine H5N1 in C57BL/6J mice (18, 21), arguing that the observed difference may be mouse strain dependent rather than inoculation route dependent.

A previous study showed that H5N1’s tropism for the nasal turbinate in BALB/c mice is influenced by the amino acid at position 627 in PB2: a lysine (K) at this position promotes replication in the nasal turbinates and lungs, while a glutamic acid (E) restricts it (19). Likewise, a secondary study showed the selection of 627K within HPAI H5N1 in the respiratory tract of a mouse model, and an increase in virus replication in the upper and lower respiratory tract associated with this mutation (17). In our study, VN1203, which contains the amino acid PB2 627K, displays restricted virus replication in the nasal turbinates compared to Bov342, but replicates effectively in lung tissue. Other animal models also do not show a clear difference in virus replication based on the PB2 E627K mutation. In ferrets, no differences in shedding from the upper respiratory tract was noted between reverse genetics viruses only differing by the PB2 627 amino acid, and PB2 627E was stable in a ferret serial passage study (27). In guinea pigs, no major differences were found in transmission of VN1203 with PB2 627K or 627E. Additionally, introduction of the PB2 627E mutation in a H3N2 isolate only resulted in moderate differences in virus replication in the upper and lower respiratory tract compared to H3N2 containing PB2 627K (20). Thus, tissue tropism may vary depending on the animal model used.

HPAI H5Nx infection differs from other influenza A viruses as it has a significantly higher likelihood of leading to CNS infection within mammalian hosts (Reviewed in (28)). For example, experimental infection of cats via intratracheal inoculation or feeding on H5N1-infected chicks resulted in infection of multiple regions of the brain (29). In contrast, CNS involvement has not yet been reported for dairy cows, either via natural or experimental infection. It has been hypothesized that H5N1 transmission within and between cattle herds is associated with milking practices. HPAI H5N1 replication was limited to the mammary glands when lactating cows were inoculated via the mammary gland (30). Thus, CNS involvement may be associated with efficient replication in the respiratory tract. In our study, CNS involvement of VN1203-inoculated animals was limited, and so was replication the upper respiratory tract.

Unsurprisingly, we detected significant differences in cytokine and chemokine profiles in brain and nasal turbinate tissues when comparing Bov342 and VN1203 at endpoint: induction of cytokines and chemokines was mostly limited to tissues obtained from animals exposed to Bov342 in which extensive virus replication was found. However, despite similar virus replication in lung tissue, significantly different cytokine and chemokine profiles were found between Bov342 and VN1203-exposed animals at protein level. Bov342 infection resulted in upregulation of CXCL1, whereas VN1203 resulted in IFN-γ, IL-2, IL-5, and IL-10. At the RNA level, gene expression was relatively similar between isolates. PCA showed clustering of the majority of lung samples, independent of virus isolate exposure, and histologic examination of the lungs did not reveal any striking differences between Bov342 and VN1203, both isolates caused a similar necrotizing bronchiolitis to bronchointerstitial pneumonia. It should be noted that differences may be time-related, Bov342-exposed animals reached endpoint at 5 days post exposure, and VN1203-exposed animals reached endpoint criteria at 6 days post exposure. More detailed analyses would be needed to confirm these findings and identify mechanisms of induction.

The key histologic differences between VN1203 and Bov342 exposure was observed in the olfactory neuroepithelium and CNS. By examining sagittal sections of the whole head in these H5N1-exposed animals, we were able to visualize the distribution of viral antigen in the nasal turbinates and brain in relation to one another. In the Bov342 animals, there was extensive virus antigen observed in the olfactory neuroepithelium, olfactory bulb, and scattered throughout other regions of the brain. Given the consistent and abundant immunoreactivity in the olfactory tissues, extension of the virus from the olfactory neuroepithelium through the nerves to the olfactory bulb seems most likely. This is supported by occasional NP antigen being observed in olfactory nerves. However, the presence of NP antigen in random foci throughout the CNS, distant from the olfactory bulb, and other tissues in the head suggest hematogenous dissemination might also plays a role. While histopathologic lesions were not observed in these areas of abundant immunoreactivity, the alterations in cytokine profiles in these animals compared to the VN1203 group indicates a cellular response. As these animals were euthanized prior to the onset of severe neurologic disease, it may be that histologic lesions would develop later. A publication describing the examination of cats who had died on dairy farms, and later tested positive for H5N1, had a severe meningoencephalitis and chorioretinitis (11). Interestingly, viral antigen was found not only in the CNS Bov342-exposed animals, but also within the retina.

The potential consequences of viral antigen in various tissues not typically associated with influenza A virus replication is unknown. To the author’s knowledge, influenza A antigen has not been documented in some of these sites, which could either be a function of not assessing these sites on routine sample collection or uniquely wide tropism of Bov342 in this mouse model. While we did not observe any pathology in these unexpected sites of virus antigen, further investigation into potential sequalae is warranted. Of particular interest to the authors is the presences of viral antigen in the choroid plexus, where virus may be shed into the cerebral spinal fluid. Additional animal studies evaluating neurotropism of Bov342 should consider evaluating the CSF and spinal cord for viral load and histology.

In conclusion, we show efficient virus replication of H5N1 strains VN1203 and Bov342 in C57BL/6J mice upon aerosol exposure. In this study, Bov342 displays a neurotropic profile, whereas both viruses replicate efficiently in the lower respiratory tract. Surprisingly, differing cytokine profiles can be detected in lung tissue at endpoint, dependent on virus isolate. Importantly, the findings in this study appear to be mouse specific and likely not representative of what will occur in humans, as thus far there is no evidence of CNS infection in contemporary H5N1 patients. Understanding the strain differences in the context of the mouse model is still of great value and these models will be essential for the evaluation of vaccines and antivirals.

## Methods

### Ethics statement

Mouse studies were performed in an AAALAC International-accredited facility and approved by the Rocky Mountain Laboratories Institutional Care and Use Committee following the guidelines put forth in the Guide for the Care and Use of Laboratory Animals 8th edition, the Animal Welfare Act, United States Department of Agriculture and the United States Public Health Service Policy on the Humane Care and Use of Laboratory Animals.

The Institutional Biosafety Committee (IBC) approved work with highly pathogenic H5N1 avian influenza viruses under BSL3 conditions. Virus inactivation of all samples was performed according to IBC-approved standard operating procedures for the removal of specimens from high containment areas.

### Cells and Viruses

Bovine HPAI H5N1 isolate A/Bovine/Ohio/B24osu-342/2024 (EPI_ISL_678615) was obtained from Richard Webby at St. Jude’s Children hospital, Memphis, TN, USA and Andrew Bowman at Ohio State University, Columbus, OH, USA. Human HPAI H5N1 isolate A/Vietnam/1203/2004 was obtained from Dr. Kanta Subbarao at the Peter Doherty Institute, Melbourne, Australia.

Virus propagation was performed in Madin-Darby canine kidney (MDCK) cells in MEM supplemented with 1 mM L-glutamine, 50 U/mL penicillin, 50 μg/mL streptomycin, 1x NEAA, 20 mM HEPES, and 4 µg/mL TPCK trypsin. MDCK cells were maintained in MEM supplemented with 10% fetal bovine serum, 1 mM L-glutamine, 50 U/ml penicillin, and 50 μg/ml streptomycin, 1x NEAA, 20 mM HEPES. Mycoplasma testing is performed at regular intervals. No mycoplasma was detected during the study.

### Animal studies

Six groups of C57BL/6J mice (Jackson Laboratories, strain #000664, between 6-8 weeks old) were challenged with HPAI H5N1 via aerosols (see details below), resulting in an estimated total dose of 2.4 x 10^2^, 5.4 x 10^2^, or 1.2 x 10^4^ TCID_50_/animal for A/Bovine/Ohio/B24osu-342/2024, and 2.5 x 10^2^, 7.8 x 10^2^, or 3.6 x 10^3^ TCID_50_/animal for A/Vietnam/1203/2004. Prior to challenge, a subgroup of mice was implanted with telemetry transponders (UCT-2112, Unified Information Devices (UID)) via subcutaneous implantation and allowed to rest for three days before challenge. Activity (total inches travelled) was recorded using a UID telemetry system and UID Mouse Matrix Plates. Data was recorded continuously with a zone interval of 250 ms, 2 cycles per series, and a 1s series delay. Oropharyngeal swabs were collected every other day in 1 mL of DMEM containing 2% of FBS, 1 mM L-glutamine, 50 U/mL penicillin, and 50 μg/mL streptomycin up until 10 days post exposure. On day 3 and when endpoint criteria were reached for at least one animal per group, four animals from each group were euthanized.

The lungs were excised and weighed, and samples were taken from lung, brain, and nasal turbinate tissues for virus titrations and histopathology. Six animals in each group were monitored at least daily for signs of disease.

### Virus exposure via aerosols

Aerosol droplet nuclei were generated by a three-jet collision nebulizer (Biaera Technologies), ranging from 1 µm to 5 µm in size. A sample of 6 L of air per min was collected during the 10 min exposure on the 47 mm gelatin filter (Sartorius). Post-exposure, the filters were dissolved in 10 mL of DMEM containing 10% FBS and infectious virus was titrated, and the aerosol concentration was calculated. The estimated inhaled inoculum was calculated using the respiratory minute volume rates of the animals determined using methods used previously (31). The inhaled dose was calculated using the simplified formula D = R × Caero × Texp (32), where D is the inhaled dose, R is the respiratory minute volume (l min−1), Caero is the aerosol concentration (tissue culture infectious dose, TCID50) and Texp is the duration of the exposure (min). Three different groups were exposed to BoV342 and VN1203 HPAI H5N1 variants. The average doses per BoV342 group are as follows: 2.4 x 10^2^, 5.4 x 10^2^, and 1.2 x 10^4^ TCID_50_. The average doses per VN1203 group are as follows: 2.5 x 10^2^, 7.8 x 10^2^, and 3.6 x 10^3^ TCID_50_.

### Virus titration

Tissue sections were weighed and homogenized in 1 mL of MEM. Virus titrations were performed by end-point titration of 10-fold dilutions of virus swab media or tissue homogenates on MDCK cells in 96- well plates. When titrating tissue homogenate, the top 2 rows of cells were washed two times with PBS prior to the addition of a final 100 µl of MEM supplemented with 1 mM L-glutamine, 50 U/mL penicillin, 50 μg/mL streptomycin, 1x NEAA, 20 mM HEPES, and 4 µg/mL TPCK trypsin. Cells were incubated at 37 °C and 5% CO2. After three days, presence or absence of infectious virus was measured using a HA assay. Turkey red blood cells (IGTKWBNAC50ML, Innovative Research) were ordered weekly. Red blood cells (RBCs) were washed at least three times with PBS and diluted to 0.33% in PBS directly before use. 75 µl of 0.33% RBCs were added to 25 µl of virus and left at 4°C for 1 hour. Hereafter, wells were marked as either agglutinated or negative. Titers were calculated using the Spearman-Karber method.

### RNA extraction

Tissue samples were bead homogenized and extracted using the RNeasy kit (Qiagen) according to the manufacturer’s instructions and following high containment laboratory protocols.

### Quantitative reverse-transcription polymerase chain reaction

Following extraction, 17 µL of RNA from nasal turbinates, lung, and brain samples were treated with TURBO DNase (Invitrogen) according to the manufacturer’s instructions. After DNase treatment, the RNA was diluted 1:5 in molecular grade water (Invitrogen). DNase-treated RNA (5 µL) was used for each host gene assessed (**Supplementary Table 1**). Fold change was calculated for host genes by dividing the sample relative gene expression by the average relative gene expression of the healthy controls (ΔΔCt) and then log_2_ transformed.

### Measurement of cytokines and chemokines protein levels in tissues

Tissue samples were bead homogenized in 1 mL of MEM and spun for 10 min at 8000 x rcf. 150 µL of supernatant was inactivated with γ-irradiation (8 MRad) according to standard operating procedures.

Tissue supernatants were assayed using the MSD Technology V-PLEX Proinflammatory Panel 1 Mouse kit (K-15048D, includes CXCL1, IFN-γ, IL-1β, IL-2, IL-4, IL-5, IL-6, IL-10, IL-12p70, and TNFα) according to the manufacturer’s instructions. Analysis was performed using Discovery Workbench LSR_4_0_13 Software.

### Histology and immunohistochemistry

All tissues harvested at the time of necropsy were fixed in 10% neutral-buffered formalin for ≥7 days. Hematoxylin and eosin (H&E) staining, and immunohistochemistry (IHC) were performed on formalin-fixed paraffin-embedded (FFPE) tissue sections. Immunoreactivity was detected using Millipore Sigma Anti-Influenza A nucleoprotein antibody (Cat.#ABF1820-25UL) at a 1:12,000 dilution. Roche Tissue Diagnostics DISCOVERY Omnimap anti-rabbit HRP (#760-4311) was used as a secondary antibody. For negative controls, replicate sections from each control block were stained in parallel following an identical protocol, with the primary antibody replaced by Vector Laboratories rabbit IgG (#I-1000-5) at a 1:2500 dilution. The tissues were stained using the DISCOVERY ULTRA automated stainer (Ventana Medical Systems) with a Roche Tissue Diagnostics DISCOVERY purple kit (#760-229).

Histologic lesions from the brain, nasal turbinates and lung were categorized and scored by a board-certified veterinary pathologist blinded to group allocations (**Supplementary Table 2**). For each category, histological lesions were scored on the scale of 0 = none, 1 = rare, 2 = mild, 3 = moderate and 4 = severe. Immunohistochemistry scoring was as follows 0 = none/no positive cells; 1 = rare positive cells; 2 = few positive cells; 3 = moderate numbers of positive cells; 4 = abundant positive cells. A representative lesion from each group was selected for figures.

### Enzyme-linked immunosorbent assay

Nunc MaxiSorp flat bottom 96-well plates (ThermoFisher Scientific) were coated with 50 ng in 50 µl/well of influenza A H5N1 A/Vietnam/1203/2004 hemagglutinin (HA) protein (IBT Bioservices, 1501- 001), or A/bald eagle/Florida/W22-134-OP/2022 neuraminidase (NA) protein (BEI Resources, NR- 59476), blocked with 100 µL of casein in PBS (ThermoFisher Scientific), and incubated with serially diluted mouse sera (1:100-1:512000) in duplicate. Immunoglobulin G (IgG) antibodies were detected by using affinity-purified polyclonal antibody peroxidase-labeled donkey-anti mice IgG (Jackson Immunoresearch, 715-035-151) or affinity-purified polyclonal antibody peroxidase-labeled goat anti-monkey IgG (ThermoFisher Scientific, PA1-84631) at a dilution of 1:5000 in casein followed by 3,3′,5,5′- Tetramethylbenzidine 2-component peroxidase substrate (Seracare, 5120–0047) and stop solution (Seracare, 5150-0021). The optical density at 450 nm (OD450) was then measured. Wells were washed three times with PBS containing 0.1% Tween in between each step.

### Statistical Analysis

Principal Component Analysis (PCA) was run at day 3 and endpoint, and included viral titer, protein abundances, and mRNA levels in brain, lung, and nasal turbinate samples with all data log_10_ transformed. Variables with more than 10% missingness (IFNβ and IFN-l2 mRNA levels) or with low variability (IL12p70, IL-4, and IL-2 protein abundance) were dropped from PCA analyses, and the remaining missing observations were imputed using Multiple Imputation by Chained Equations (MICE) (33). The number of clusters for each PCA were determined by the majority vote of 26 commonly used indices for identifying the optimal number of clusters under k-means clustering (34), and k-means clusters were calculated for each PCA. PCA analyses and visualizations were run in R version 4.4.1.

All other statistical analysis were performed in Graphpad Prism using a two-way ANOVA followed by a Mann-Whitney test. Gene expression differences between viruses and dose were compared at day 3 and endpoint using a two-way ANOVA with Tukey’s post-test, and differences across time points were assess with a two-way ANOVA with Sidak’s multiple comparisons test. A Bonferroni correction was applied, and only p-values greater than 0.0044 were considered significant.

## Data Availability Statement

All data have been deposited on Figshare: 10.6084/m9.figshare.27679509. All code has been deposited on Github: https://github.com/IDSS-NIAID/2024-H5N1-comparisons-in-mice.

## Acknowledgements

We thank the animal caretakers of the Rocky Mountain Veterinary Branch, NIAID, NIH for their assistance during the study, Dean Follmann for advice on appropriate statistical analysis, Andrea Marzi and Kyle O’Donnell for providing controls and reagents for the ELISA assay, and Anita Mora for assistance with figures.

---

**Supplementary Table 1.**
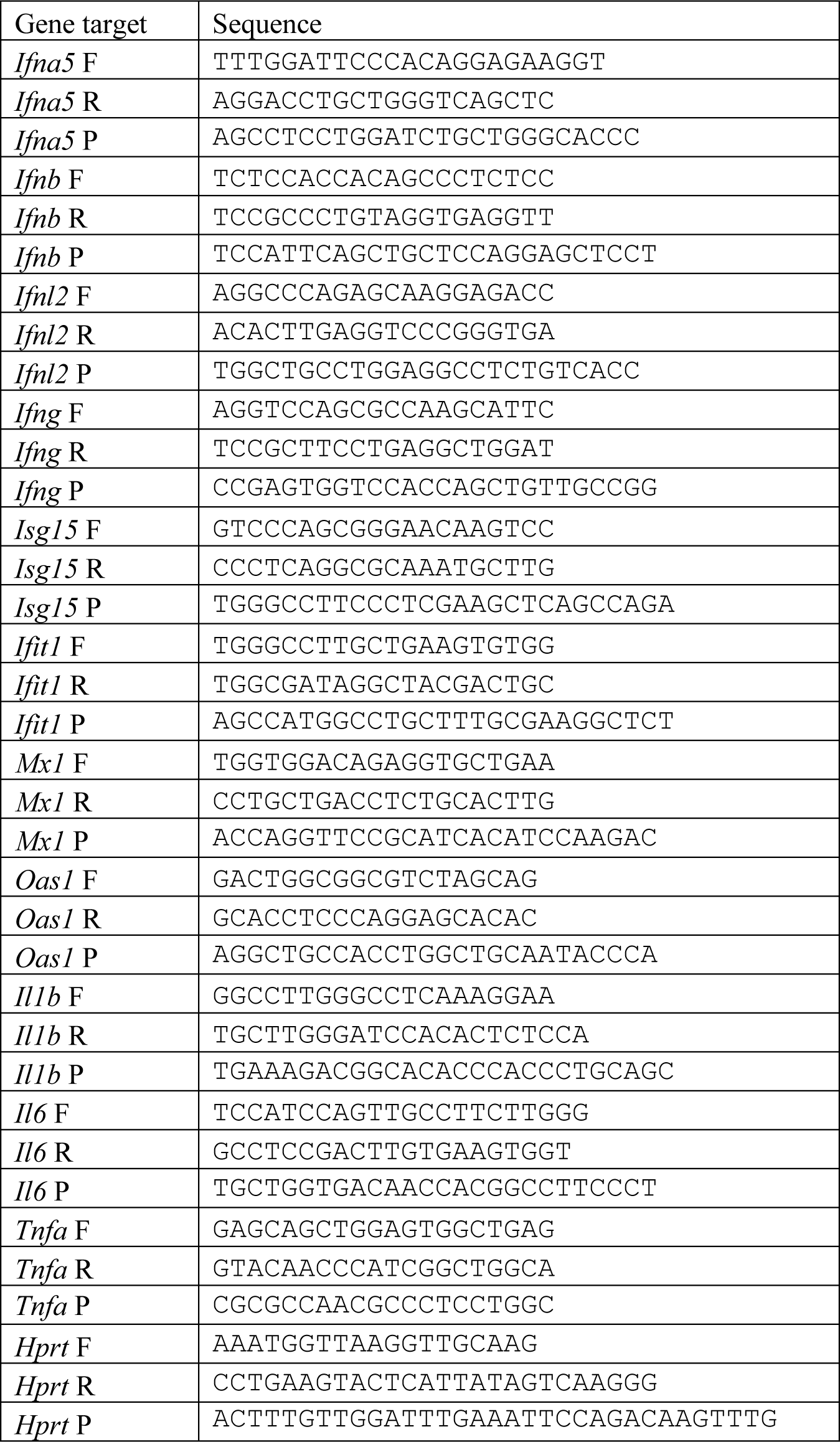
RT-qPCR primers used to measure host gene expression.

**Supplementary Table 2:**
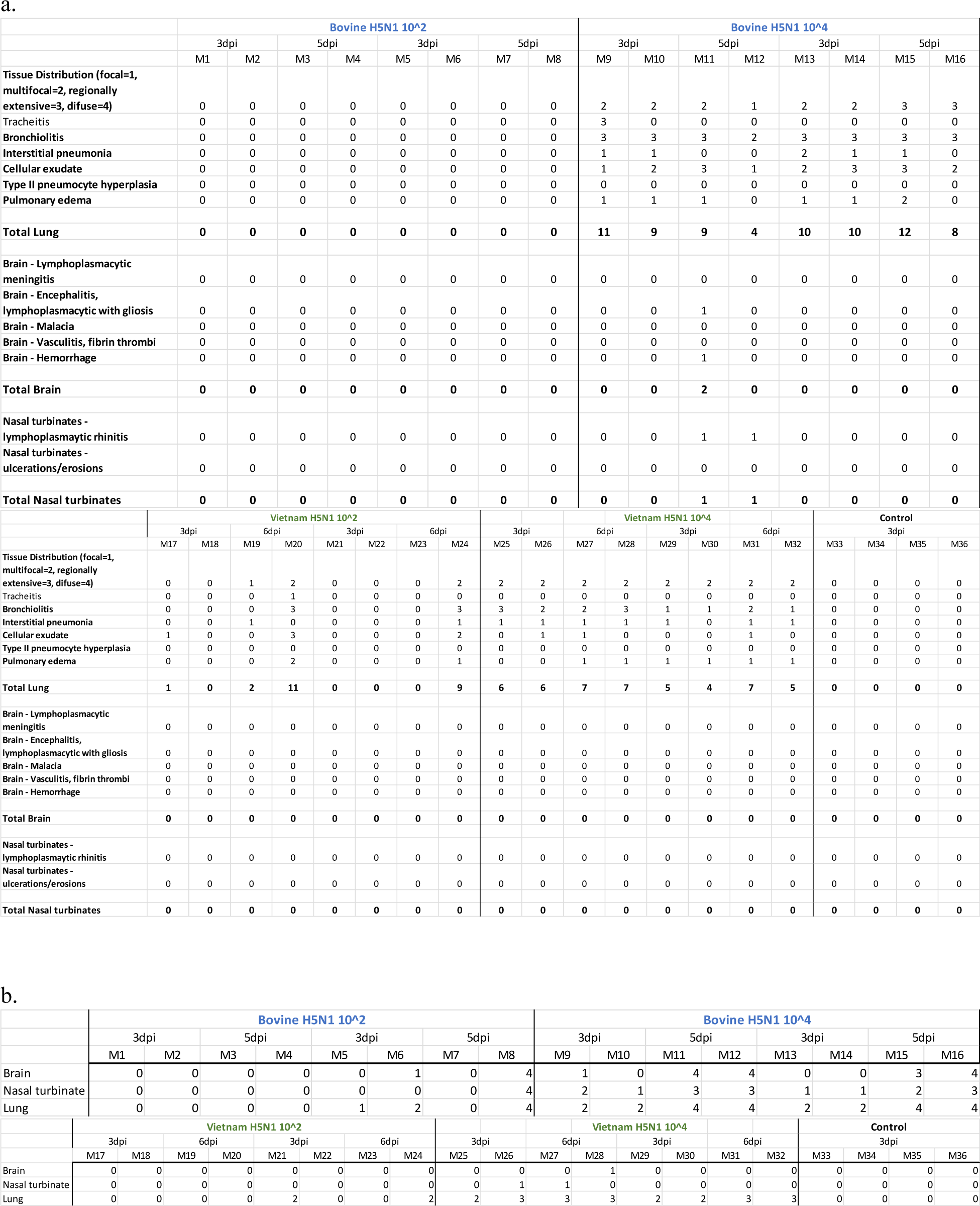
Semiquantitative scores assigned by a board-certified veterinary pathologist following histopathologic examination of H&Es (a) and immunohistochemical labeling for influenza A virus NP (b) of the lungs, brains, and nasal turbinates.

**Supplementary Figure 1.**
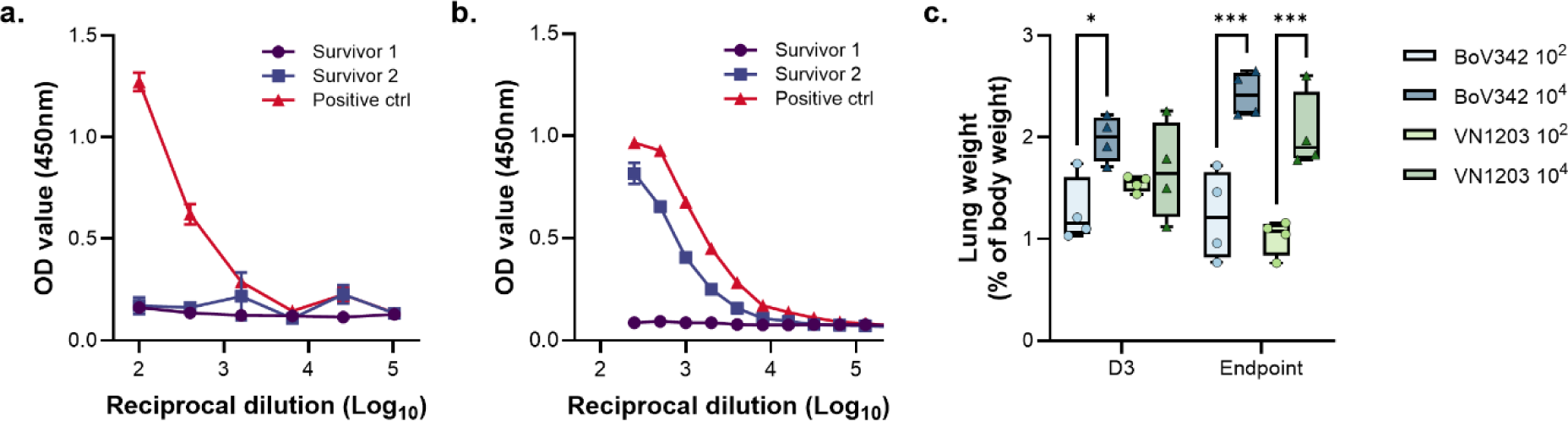
a. Binding antibody titers to HA of VN1203 of two mice that survived VN1203 challenge compared to positive control mouse sera. b. Binding antibody titers to NA of A/bald eagle/Florida/W22-134-OP/2022 of two mice that survived VN1203 challenge compared to positive control NHP sera. c. Lung body weight ratio of animals inoculated with Bov342 or VN1203. Statistical significance was determined via an ordinary two-way ANOVA followed by Tukey’s multiple comparisons test. * = p-value <0.05; *** = p-value <0.001.

**Supplementary Figure 2.**
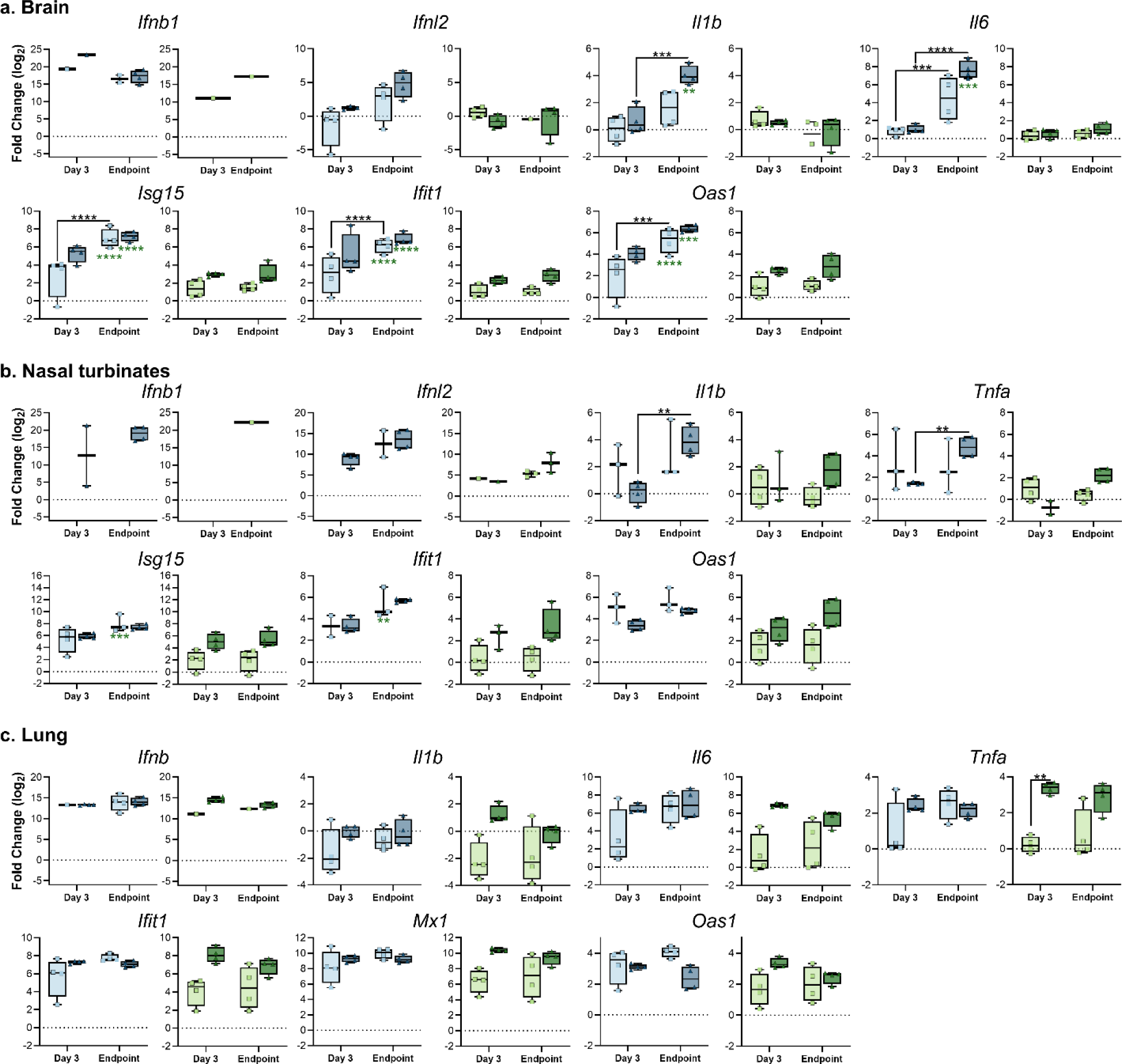
Aerosol inoculation of C57Bl/6 mice with H5N1 results in virus-, tissue- and dose-dependent induction of cytokines and antiviral effector genes. Gene expression in brain (a), nasal turbinate (b), or lung (c) shown as log_2_ transformed fold change relative to healthy control mice. Individual values showed, bar graph shows minimum to maximum, with middle line displaying median (N=4). For genes with cycle threshold values below the limit of detection, no value is reported. Gene expression differences between viruses and dose were compared at day 3 and endpoint using a two-way ANOVA with Tukey’s post-test, and differences across time points were assess with a two-way ANOVA with Sidak’s multiple comparisons test. A Bonferroni correction was applied, and only p-values greater than 0.0044 were considered significant. **= p-value <0.0044, ***= p-value <0.001, ****=p-value <0.0001.

**Supplementary Figure 3.**
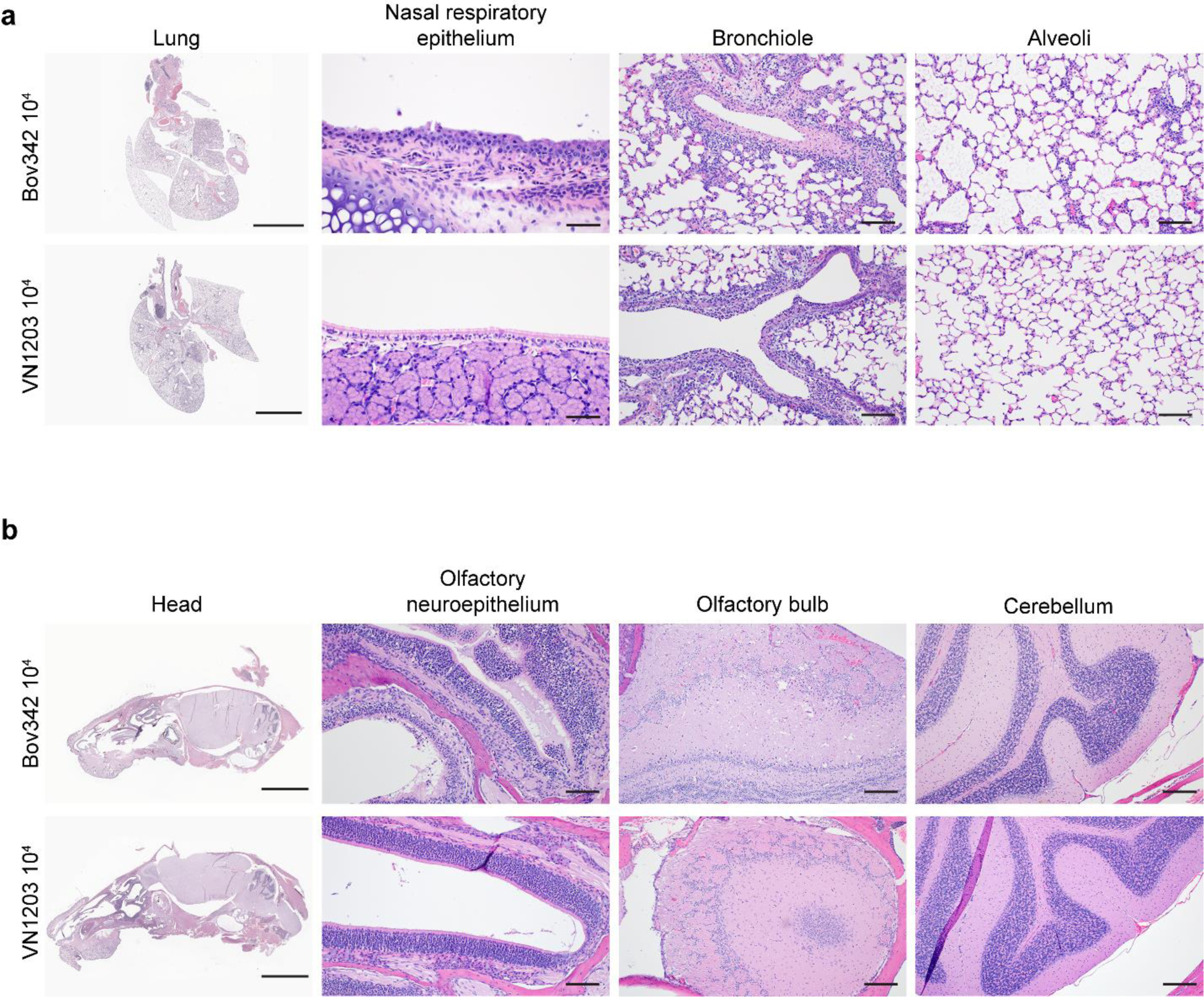
Aerosol inoculation of C57BL/6J mice with 10^4^ TCID50 of both Bov342 and VN1203 result in necrotizing bronchiolitis, and no observable pathology in the CNS. Hematoxylin and eosin (H&E) staining of lungs (a) and head (b). In the lungs, the primary histopathologic finding was a necrotizing bronchiolitis (a). Minimal pathology was observed in the nasal respiratory epithelium, and an occasional interstitial pneumonia was observed surrounding affected bronchioles (a). Significant pathology was not observed in the olfactory neuroepithelium or CNS in either Bov342 or VN1203 groups. All images were selected from representative animals in the endpoint groups. a. Lung: scale bar=4mm. Nasal respiratory epithelium: scale bar=50µm. Bronchiole: scale bar=100µm. Alveoli: scale bar=100µm. b. Head: scale bar=5mm. Olfactory neuroepithelium: scale bar=100µm. Olfactory bulb: scale bar=200µm. Cerebellum: scale bar=200 µm.

**Supplementary Figure 4.**
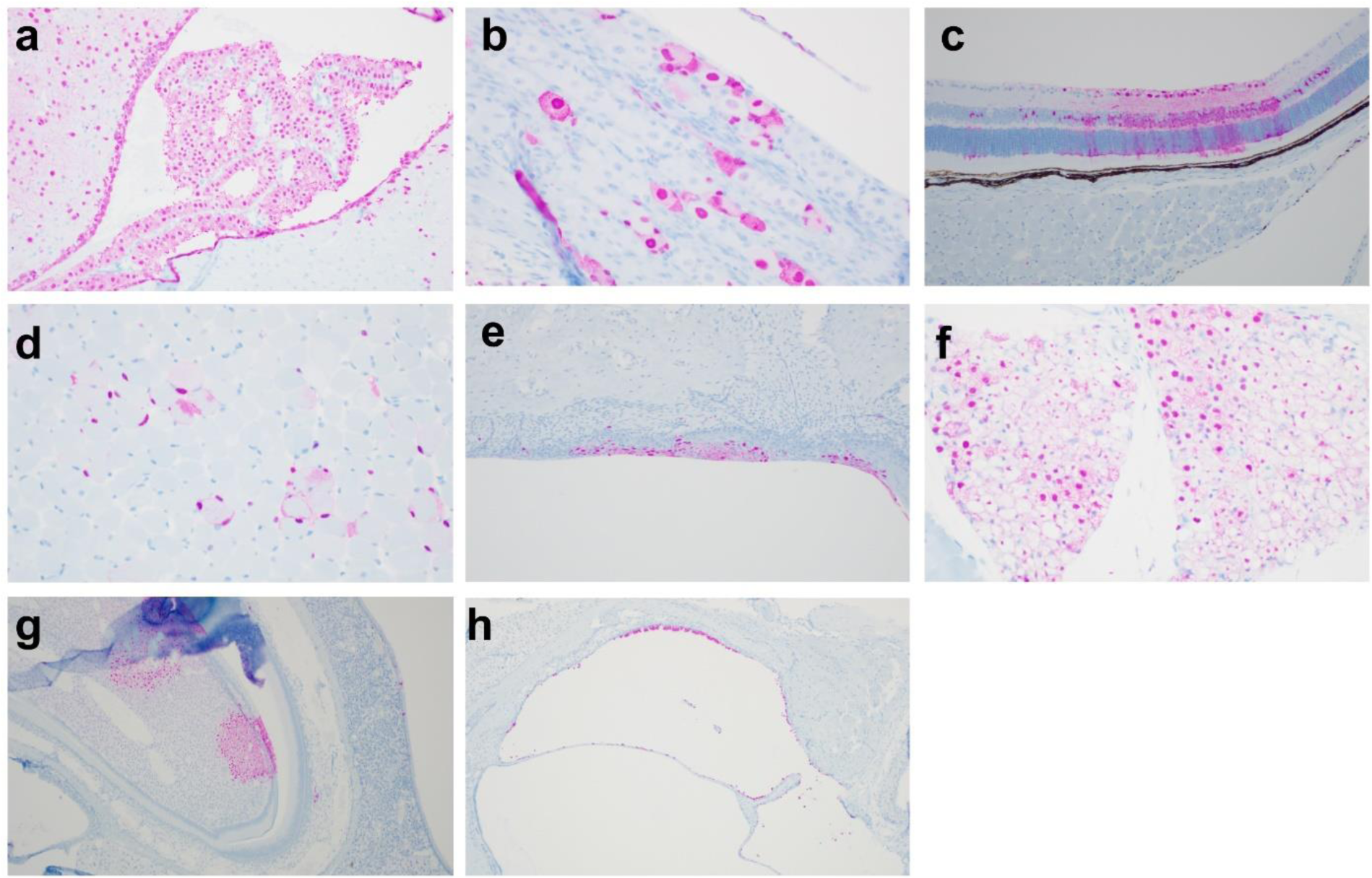
**Immunoreactivity for Influenza A NP was observed in multiple atypical sites within the head**. IAV NP immunoreactivity (pink) was observed within the choroid plexus and ependymal cells lining the ventricle (a), large neuronal cells composing a nuclei likely associated with a cranial nerve (b), multiple cell layers of the retina (c), skeletal myocytes (d), mucosal epithelium overlying the hard palate (e), within brown adipocytes (f), within a tooth root and the overlying odontoblasts (g), and epithelial cells lining the inner ear (h).

**Supplementary Figure 5.**
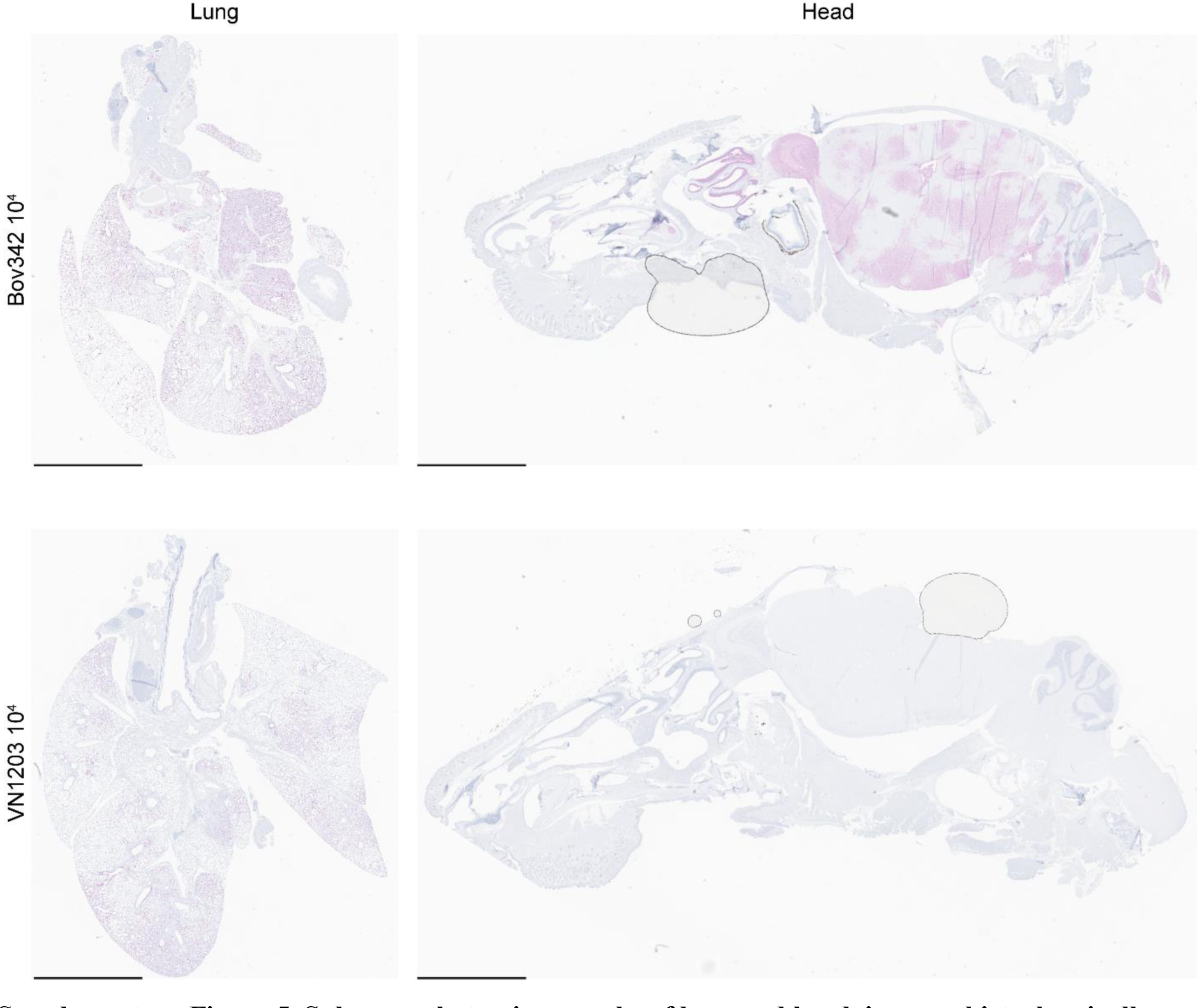
Subgross photomicrographs of lung and head, immunohistochemically labeled with Influenza A NP (pink) from animals in the endpoint groups for both isolates. Immunoreactivity was observed in the lungs of both Bov342 and VN1203 animals, however IAV NP labeling was only observed in the CNS of the Bov342 animals. Representative images selected from animals in the endpoint groups for both isolates. Lung: scale bar=4mm. Head: scale bar=5mm.

